# Positive allosteric modulators of lecithin:cholesterol acyltransferase adjust the orientation of the membrane-binding domain and alter its spatial free energy profile

**DOI:** 10.1101/2020.10.20.344978

**Authors:** Akseli Niemelä, Artturi Koivuniemi

## Abstract

Lecithin:cholesterol acyltransferase protein (LCAT) promotes the esterification reaction between cholesterol and phospholipid derived acyl chains. Positive allosteric modulators have been developed to treat LCAT deficiencies and, plausibly, also cardiovascular diseases in the future. The mechanism of action of these compounds is poorly understood. Here computational docking and atomistic molecular dynamics simulations were utilized to study the interactions between LCAT and the activating compounds. Results indicate that all drugs bind to the allosteric binding pocket in the membrane-binding domain in a similar fashion. The presence of the compounds in the allosteric site results in a distinct spatial orientation and sampling of the membrane-binding domain (MBD). The MBD’s different spatial arrangement plausibly affects the lid’s movement from closed to open state and *vice versa*, as suggested by steered molecular dynamics simulations.

## INTRODUCTION

Lecithin:cholesterol acyltransferase (LCAT) is an enzyme that is responsible for producing cholesterol esters (CEs) in circulation by linking the acyl chains of phospholipids to cholesterol (CHOL) molecules. LCAT mediated CE formation predominantly occurs in high-density lipoprotein (HDL) particles in which apolipoprotein A-I (apoA-I) serves as a cofactor for the reaction. After the formation of CEs, the morphology of HDL particles transforms from discoidal to spherical by the creation of a non-polar CE phase, which is shielded from aqueous surroundings by an amphiphilic monolayer comprised of phospholipids, CHOL, and apolipoproteins. Owing to this, the action of LCAT is an essential step in HDL mediated reverse cholesterol transport (RCT), in which intracellular CHOL and phospholipids are transported from extrahepatic tissues to the liver. Importantly, HDL particles also remove CHOL from lipid droplet-laden macrophages that are the hallmark of early atherosclerotic lesions in the arterial intima (1). For this reason, chiefly, RCT’s efficiency is hypothesized to be associated with the progression of coronary heart disease (CHD (2–4).

The normal functioning of LCAT is reduced or completely lacking in individuals suffering from the autosomal recessive disorders familial LCAT deficiency (FLD) and fish-eye disease (FED) (5–7). The clinical manifestations of LCAT deficiency include diffuse corneal opacities, target cell hemolytic anemia, and kidney failure. While the role of LCAT deficiencies and activity in CHD progression is currently far from understood, multiple ongoing research programs are attempting to shed light on LCAT activity’s boosting therapies in this context. These therapeutic approaches include recombinant human LCAT (rhLCAT) injections, biologics such as peptides and antibodies, and small molecular activators (8–11). The rhLCAT (MEDI6012, formerly ACP-501) is particularly of note as during recent phase II trials, it was shown not to cause any serious adverse events in the 48 participants, and its administration led to a dose-dependent increase in HDL-cholesterol (HDL-C) concentration (12). Intriguingly, a recent HDL particle lipidomics study in which a subsequently defined non-equilibrium reaction quotient describing global CHOL homeostasis in circulation was established suggests a deficient conversion of CHOL to CE in CHD patients when compared to controls (13). Notably, the deficient CHOL to CE conversion was determined without utilizing LCAT activity assays or plasma concentration measurements. Hence, this approach might provide a more reliable means to assess LCAT mediated CHOL to CE conversion that considers the effect of native lipid compositions better than exogenous plasma assays that are based on, e.g., the addition of labeled CHOL molecules into the plasma of individuals with differing amount of CHOL in different lipoprotein pools.

Because of the reasons above, it would be valuable to develop LCAT activity promoting therapeutics, firstly, for the treatment of different LCAT deficiencies and, secondly, for providing additional ways to elucidate the impact of LCAT based therapeutics on the functional quality of HDL particles in the context of RCT and CHD. In this respect, a set of promising LCAT activity promoting compounds was developed by Daiichi Sankyo (14–17). Recently, out of this set of compounds, the piperidinyl-pyrazolopyridine derivatives (compounds 1, 2, 3, 6, 8, and 9) were further investigated by Manthei et al., showing that the compounds increase the activity of LCAT up to 3.7-fold (18). In the same study, an X-ray structure for LCAT was solved, with both bound activator compound 1b and inhibitor isopropyl dodecyl fluorophosphonate (IDFP), revealing that compound 1b resides in the cleft of the membrane-binding domain (MBD, see Fig 1). Although the MBD has been shown to interact with lipids when LCAT is bound to lipid bilayers or HDL particles (19–22), it became evident based on binding studies that the activators do not alter the association of LCAT with discoidal HDL particles (18). In addition, the X-ray structure revealed that compound 1b forms hydrogen bonds with MET49, TYR51, ASP63, and ASN78 and possesses the enantiomeric R state (compound 1b in Fig 2).

**Figure 1.**
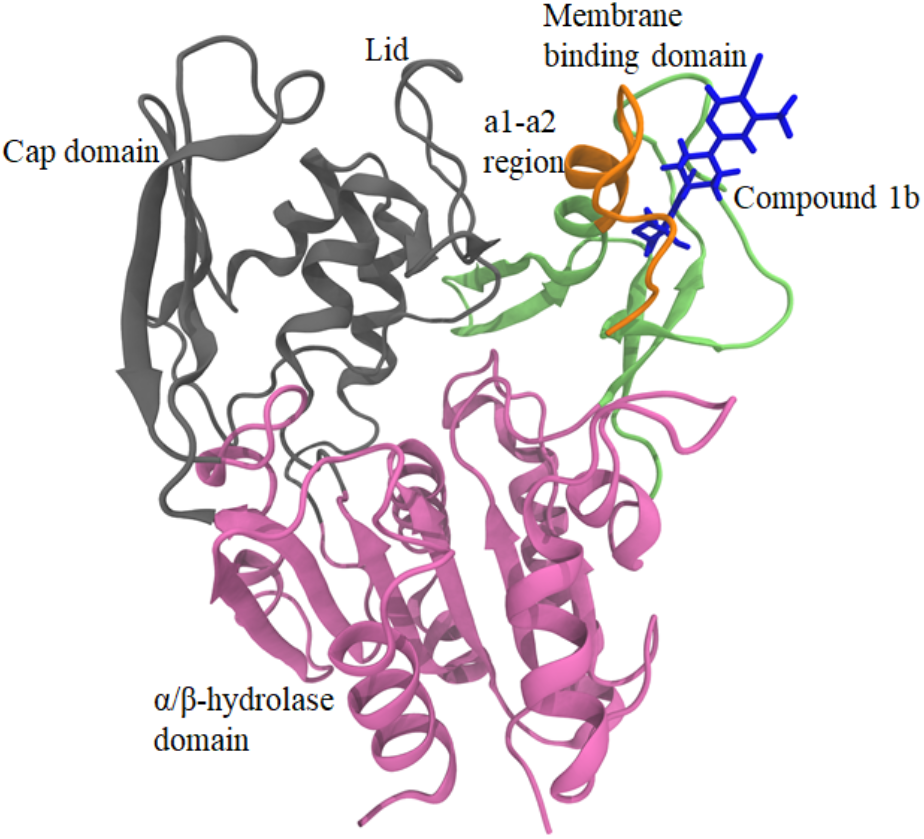
The structure of LCAT with compound 1b bound to the allosteric site. The α/β hydrolase domain is colored as purple, the cap domain as gray, and the membrane-binding domain as green. The a1-a2 region of the MBD (amino acids 63-75) is marked with orange. The protein is rendered as a cartoon secondary structure representation and the compound as blue sticks.

**Figure 2.**
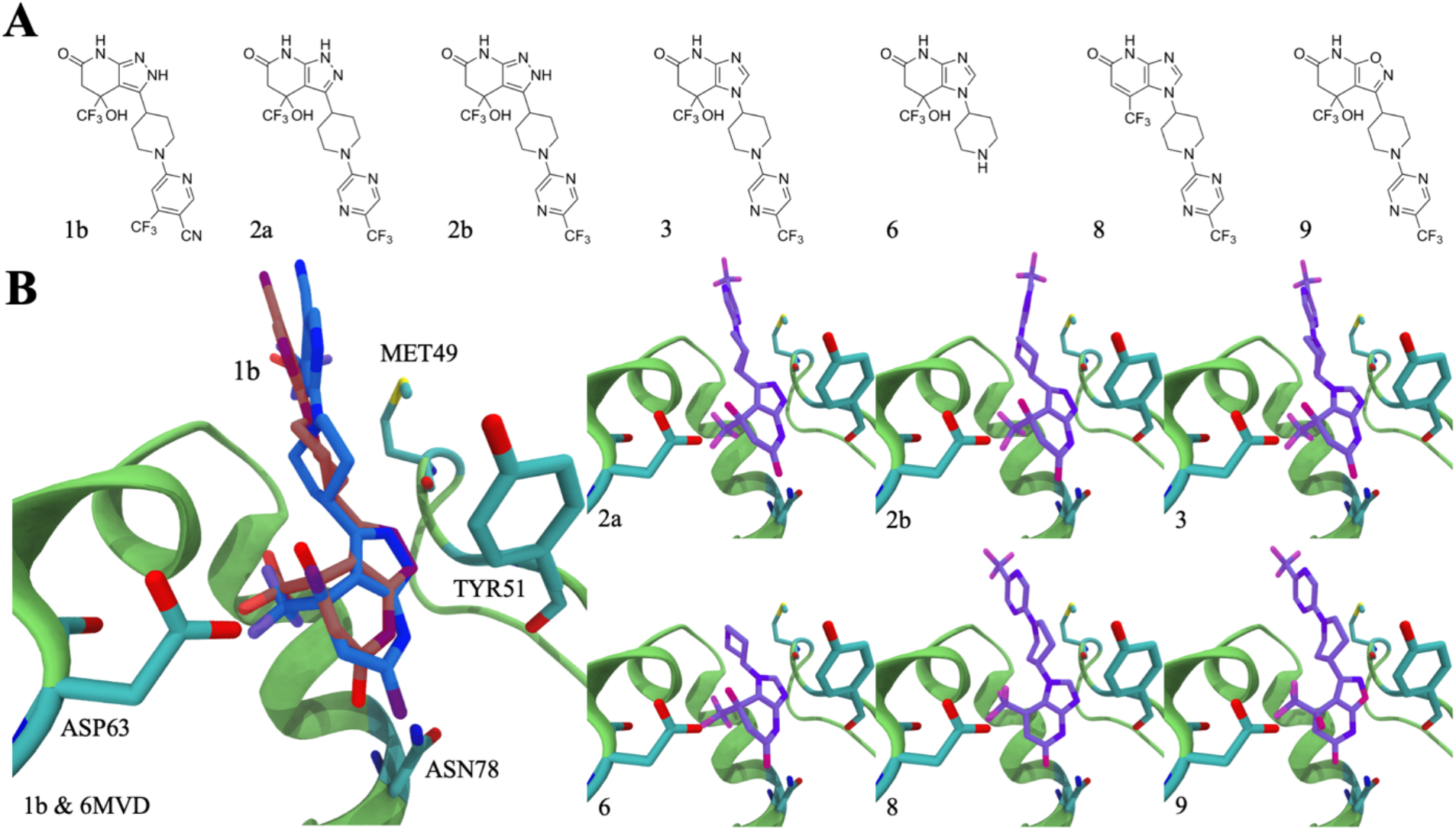
**(A)** The chemical structures of the compounds studied. **(B)** The docking poses of the compounds bound (hued purple) in the MBD cleft of LCAT and the co-crystallized compound 1b (PDB accession code 6MVD; hued blue) compared to its position docked back the structure (hued red). The allosteric site of LCAT is rendered as a cartoon secondary structure representation and shown as green. The compounds and the significant amino acids are rendered as sticks and are colored according to different elements. Carbon atoms are cyan, oxygen red, nitrogen blue, fluorine pink, and sulfur yellow.

Interestingly, previous hydrogen-deuterium exchange (HDX) studies indicated that the α-helical folds a1 and a2 (comprised of amino acids 63-75) in the MBD and the lid loop region are more dynamic when compared to the rest of the LCAT structure (Fig 1). It was further demonstrated that the binding of IDFP decreased HDX in the a1-a2 region. While the X-ray structure of LCAT with bound compound 1b and IDFP also indicated that the temperature factors are slightly decreased in the a1-a2 site when compared with LCAT structures without drugs, it is impossible to determine what is the impact of compound 1b alone on the dynamics of the a1-a2 region and the whole MBD in general. Besides, the constraints imposed by different crystal lattices might not reveal real conformational preferences for the local structural domains of enzymes compared to conditions in which the enzymes are in fully aqueous surroundings and under the influence of the Brownian motion. Significantly, this might be the case when highly mobile regions of proteins are considered. Nevertheless, the binding of IDFP alone might induce the opening of the lid as was suggested by the HDX data (23) and, therefore, the conformation of the lid loop in the X-ray structure of LCAT with bound compound 1b might not resemble the case when compound 1b is solely present. However, as compound 1b forms hydrogen bonds with the two amino acids located in the a1-a2 region, namely with ASP63 and ASN78, it might have an additional effect on, e.g., the rigidity of the region that may play an essential role in the activation of LCAT.

Consequently, our aim in this study was to computationally characterize how the set of positive allosteric modulators previously investigated by Manthei et al. (18) interact with the MBD and if they alter its conformation and dynamics. This, particularly, without the presence of the inhibitor IDFP. Firstly, we utilized molecular docking calculations and atomistic molecular dynamics simulations to show that the compounds interact with LCAT through a varying number and network of hydrogen bonds, which presumably stabilizes the MBD. Secondly, simulations highlight that the drugs adjust the MBD’s spatial orientation with respect to the lid in its open state. Meanwhile, accelerated weight histogram-based free energy simulations demonstrate that the compounds modify the MBD’s energetic landscape.

Further, steered molecular dynamics simulations reveal that enforcing the lid loop to a close state is accompanied by a retraction of the MBD away from the lid binding cavity. To conclude, our findings suggest that the reorientation and the different energetic landscape of the MBD induced by the allosteric compounds may facilitate the lid’s opening, therefore providing a plausible explanation of why the set of positive allosteric modulators studied here promote the activity of LCAT. Besides, this finding may be insightful when deciphering how apoA-I facilitates the activation of LCAT. In general, the results provided here pave the way for the design of new therapeutic approaches against LCAT deficiencies and for scrutinizing the HDL quality hypothesis in the context of RCT and CHD.

## MATERIALS AND METHODS

### Construction of the LCAT structure

The X-ray structure of LCAT with bound molecular activator compound 1b (18) was acquired from the Brookhaven databank (PDB ID code:6MVD). The missing two amino acid residues located at the lid region of LCAT were incorporated into the structure with the Modeller homology modeling package, as described earlier (19, 24). The standard parameters were used in the incorporation, and the structure with the lowest DOPE score was selected for the further modeling stages.

### Docking calculations

The molecular structures and initial coordinate files for all compounds were built with the Avogadro molecular editor and visualizer software (25), after which the compounds were docked to the membrane-binding domain of LCAT utilizing the Autodock Vina version 1.1.2 (26). The size of the docking grid was set big enough to cover the membrane-binding domain of LCAT. The number of grid points was set to 80 in the X, Y, and Z dimensions when the grid points’ distance was 0.375 Å. The exhaustiveness was set to 100. Nine docking configurations with the lowest binding free energies were produced, and the lowest one was selected as a starting point for molecular dynamics simulations.

### Force fields and parametrization of drug compounds for molecular dynamics simulation purposes

The AMBER99SB-ILDNS force field parameter set was used to describe the LCAT protein (27). Water was described by the TIP3P parameters (28). The initial structures for the drug compounds were built with the Avogadro program, after which the geometries of the compounds were optimized with the Gaussian software version 16 revision A.03 (29). Hartree-Fock method and 6-31G* basis set were used in the optimization procedure. The partial charges were derived by first determining the electrostatic potential around each molecule with the Gaussian program utilizing the Merz-Kollman scheme (30). The same method and basis set was employed in this stage as in the geometry optimization step. This was followed by the derivation of partial charges utilizing the restrained electrostatic potential (RESP) method to produce the QM derived electrostatic potentials around the molecules. The antechamber program included in the Amber18 modeling package was used for this purpose (31). The Lennard-Jones and bonded parameters for the compounds were taken from the GAFF force field (32).

### Simulated systems and simulation parameters

The structure of LCAT with and without drugs was placed into the center of a box with dimensions of 10 × 10 × 10 nm. Two replicates were run for the LCAT system without drugs. The compounds were incorporated into the allosteric binding site of the MBD domain based on the docking results. Each system was solvated, resulting in 14597 water molecules per system. Ions were added to neutralize the total charges of the systems. All simulations were coupled to a temperature bath of 310 K utilizing the v-rescale thermostat with a coupling constant of 0.5 ps (33). The 1.013 bar pressure was described as isotropically using the Parrinello-Rahman barostat with a coupling constant of 10 ps (34). To handle electrostatics, the Particle-Mesh Ewald (PME) summation scheme was employed with a real-space cut-off of 1.0 nm (35). The Lennard-Jones interaction cut-off was set to 1.0 nm. All systems were first energy minimized utilizing the steepest descent method followed by molecular dynamics simulations up to 1 us with the GROMACS simulation package (36, 37).

### Accelerated weight histogram simulations

The accelerated weight histogram method is an adaptive biasing method implemented in the GROMACS package that can be employed to calculate the potential mean force profiles as a function of reaction coordinates (36, 37). The approach flattens the free energy barriers along a reaction path by introducing potentials that elevate free energy minima resulting in unrestricted diffusion of the selected atoms or molecules along the reaction coordinate. Therefore, the system’s sampling is artificially enhanced, and the spatial free energy can be explored, unlike in non-adaptive biasing simulations (38). In this study, the AWH method (38, 39) was employed to probe the free energy profile when the MBD domain of LCAT changes its orientation with respect to the lid. The α-carbon atoms of amino acids I231 and M66 were chosen as the reference and pull groups, respectively. The reaction coordinate was defined as the distance between these two atoms, with the largest distance being 1.1 nm and the smallest 0.5 nm. The AWH potential was set to the umbrella, and the force constant and initial error for AWH calculations were set to 128000 kJ/mol/nm^2 and 5 kJ/mol, respectively. The estimated diffusion parameter of 0.0001 nm^2/ps was used for the coordinate dimension. The AWH systems were simulated up to 500 ns. The converge of free energy profiles and distribution profiles were registered to take place after 300 ns. The free energy profiles after 500 ns simulations were constructed utilizing the gmx awh program included in the GROMACS simulation package.

### Steered molecular dynamics simulations

To monitor the movement of the MBD domain of LCAT during the conformational change of the lid from the open to a closed state, we carried out steered molecular dynamics simulations during which the lid was pulled to a closed state. The center of mass α-carbon atoms of amino acids 110-130 was used as a reference group during the pulling simulation, whereas the α-carbon atom of amino acid 219 was pulled towards the center of mass of the reference group. The pulling force constant and the rate was set to 5000 kJ/mol·nm^2^ and 0.0001 nm/ps, respectively. The steered MD simulations were run up to 25 ns, after which the distance between the α-carbon of LEU239 and the reference group was approximately 1 nm. The distance of MET66 Cα atom from its initial position was monitored as a function of simulation time to reveal the retraction distance of MBD when the lid is pulled to a closed state. During the pulling, the protein’s rotational and translational movement was kept fixed by spatially restraining the reference group’s backbone atoms.

### Analysis

Hydrogen bonds and distances were calculated using the gmx hbond and gmx mindist tools, respectively, that are included in the GROMACS package. The default hydrogen bond parameters were used in determining the number of hydrogen bonds formed between the drugs and LCAT. The Visual Molecular Dynamics (VMD) program was used to visualize and render the figures (40).

## RESULTS AND DISCUSSION

### All compounds bind similarly into the allosteric cleft located in the MBD of LCAT

In order to investigate the differences between the binding poses of different compounds and to produce starting configurations for atomistic molecular dynamics simulations, all compounds were docked to the MBD of LCAT (PDB accession code: 6MVD, co-crystallized with compound 1b) utilizing the AutoDock Vina docking software (26). The chemical structures for all drug compounds are shown in Fig 2A. Firstly, compound 1b was docked back into the MBD to validate the parameters and the scoring function used in the docking. As seen in Fig 2B, the Autodock program was able to find the correct pose for compound 1b, with only small differences seen mainly in the water exposed part of the drug. It is also good to note that in the X-ray structure of LCAT bound to compound 1b (6MVD), compound 1b interacts with the neighboring LCAT enzyme in the crystal lattice. This likely also affects the orientation of the water exposed part of the compound in the allosteric site. After this, the rest of the compounds (2a, 2b, 3, 6, 8, and 9) were docked similarly to the MBD. Afterward, we docked the compounds again into the allosteric site after 1 us of atomistic simulations to see if carrying out the simulations would improve binding as far as the Autodock derived binding free energies are concerned.

As expected, the results in Fig 2B point out that the preferred orientation of molecules in the allosteric site is similar in each case. While tiny differences in the spatial arrangement are seen, this is expected since the chemical groups, and their spatial regions are also different between the molecules. We found out that 1 us simulations increased the binding free energies in all cases except in the case of compound 6 (Table 1). Namely, the binding free energy of compound 6 decreased after the simulation and is lower than those of the rest of the compounds (−32 kJ/mol vs. 38-42 kJ/mol, respectively). This finding follows the previous experimental findings showing that compound 6 does not activate LCAT since its binding; for some reason, it is abolished (18). However, our docking calculations and simulations (in the later chapter) imply that compound 6 can interact favorably with the allosteric site if it can access it. As the binding free energies change somewhat after the 1 us simulations (except for compound 6), it also suggests that the compounds and the allosteric site become better adapted to each other during the simulations. The binding free energies of the compounds do not correlate with the experimental fold activities produced with MUP esterase assays, implying that the strength of binding does not explain the different fold activities (18).

**Table 1.** The calculated binding free energies for the compounds docked either to the X-ray structure or the simulated structure of LCAT (after 1 us).

| Molecule | DG<sub>bind</sub>(X-ray) [kJ/mol] | DG<sub>bind</sub> (Sim.) [kJ/mol] |
| --- | --- | --- |
| Compound 1b | -37 | -41 |
| Compound 2a | -34 | -42 |
| Compound 2b | -39 | -40 |
| Compound 3 | -35 | -38 |
| Compound 6 | -34 | -32 |
| Compound 8 | -34 | -39 |
| Compound 9 | -36 | -41 |

### All drug compounds form a stable hydrogen bonding network in the allosteric site

To gain more insight into how the LCAT activators interact with the MBD domain of LCAT, we analyzed the average number of hydrogen bonds formed between the compounds and LCAT. In addition, we identified the critical amino acid residues taking part in the formation of the hydrogen bonds with the different functional groups of the drug compounds.

Fig 3A shows the average number of hydrogen bonds between the compounds and the different residues in the MBD domain. Based on the data, it is evident that mainly ASP63, TYR51, and ASN78 form hydrogen bonds with the compounds. However, the backbone amine of MET49 showed a stable hydrogen bond with compounds 1b and 2b throughout the simulations. In the case of ASP63 and ASN78, the side chain amine and carboxyl groups were involved in forming hydrogen bonds with the hydroxyl and carbonyl groups of the compounds, respectively (See Fig 3B). TYR51 formed hydrogen bonds through the backbone amine and carboxyl groups. MET49 utilized the backbone carboxyl oxygen to bond with the amine groups of compounds 1b and 2b. Compound 2a did not form a hydrogen bond with MET49, which can be traced to the changed amine hydrogen position, which abolishes the hydrogen bond interaction with MET49. Instead, compound 2a forms a hydrogen bond with TYR51. In addition, the amine hydrogen hopping increases the average number of hydrogen bonds formed with TYR51 above all other compounds, as seen in Fig2A. Compound 9 also forms a more significant number of hydrogen bonds with TYR51 and this, in turn, arose from the close location of two hydrogen acceptors, ring oxygen and nitrogen of compound 9, which enabled them to bond with the backbone amine of TYR51 at the same time.

**Figure 3.**
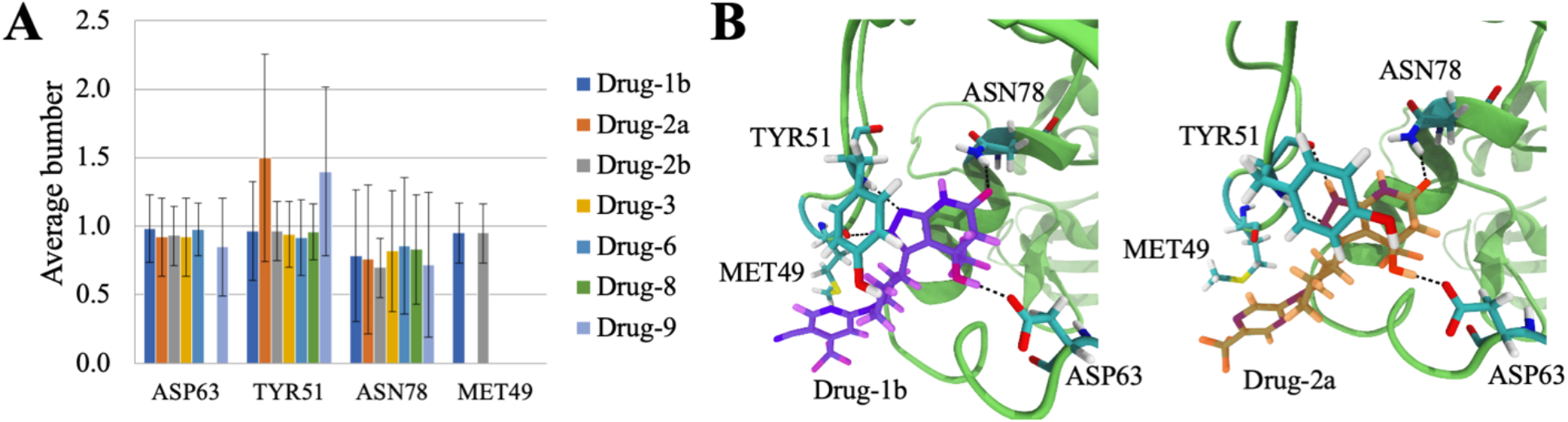
**(A)** The average number of hydrogen bonds formed between the MBD amino acids and drug compounds during the whole simulation trajectory **(B)** Visualization of the primary hydrogen bonding pairs between compounds and LCAT. Snapshots from Drug-1b (Left) and Drug-2a (Right) simulations showing the conformations of compounds 1b and 2a in the cleft of MBD. The hydrogen bonds between drugs and amino acids are marked with black dashed lines. Compounds 1b and 2a are rendered as sticks and are colored according to the element types. Compound 1b is hued purple and 2a orange. The MBD cleft is rendered as a cartoon representation and colored green. Amino acids forming hydrogen bonds with compounds are rendered as sticks and colored according to the element types. Carbon atoms are cyan, oxygen red, nitrogen blue, fluorine pink, and sulfur yellow.

Our simulations show that all drug compounds form hydrogen bonds similarly with ASP64, ASN78, and TYR51, consistent with the X-ray structure of LCAT co-crystallized with compound 1b. However, in addition to compound 1b, only compound 2b formed a hydrogen bond with MET49. Therefore, it seems that the amine group with a hydrogen donor at site 1 (Fig 3A) is required for the formation of an efficient hydrogen bond with the backbone carbonyl group of MET49. Without hydrogen at site 1, the amine group forms a hydrogen bond with the backbone amine of TYR51. The compounds studied here were partly chosen to explain the MUP esterase assay’s differences conducted by Manthei et al. 2018. Out of these, the lack of activity of compound 6 was of particular interest. As seen in our simulation analysis, compound 6 formed hydrogen bonds with the same amino acids (ASP63, TYR51, and ASN78) as the other compounds. Besides, no significant differences were detected in the average hydrogen bond numbers between compound 6 and the rest of the compounds, indicating that the hydrogen bonds are equally stable. Furthermore, according to the MUP esterase activity assays, compounds 1b, 2b, 3 had the same fold activities (2.3-2.4), whereas compounds 8 and 9 possessed the highest and lowest activation potencies (3.7 vs. 1.6), respectively. Based on our hydrogen bonding results here, we cannot argue the reason behind the different potencies.

### Positive allosteric modulator binding induces a conformational change which distances the MBD from the lid residing cavity

Next, we aimed to investigate if the compounds change the spatial arrangement of the MBD of LCAT. We calculated distances between the backbone α-carbon atoms of the MBD and the lid loop to determine if the compounds rearrange the MBD with respect to the rest of the enzyme. Upon examining the distance matrix, a conformational change was discovered where systems with drugs had the MBD pushed further from the lid loop and cavity (Fig 4A). Since the distance matrix analysis showed the average distances over the whole simulation trajectories, we examined how consistent the conformational shift was by monitoring the distance between the backbone α-atoms of M66 and I231 as a function of time (Fig 4B). It was found that the distance between the backbone α-atoms was persistently more considerable in the simulations with drug compounds bound to the allosteric site than in simulations without allosteric modulators. This conformational shift is illustrated in Fig 4C with two snapshots with and without drug molecules. To ensure that our shift is genuinely dependent on whether a compound is bound to the allosteric site or not, we removed compound 2a from the LCAT structure after an 800 ns simulation. As seen in Fig 4D, the distance between the α-atoms decreased from 0.9 to 0.7 nm in 50 ns, indicating that the MBD shifts its orientation closer to the lid cavity in the absence of the drug.

**Figure 4.**
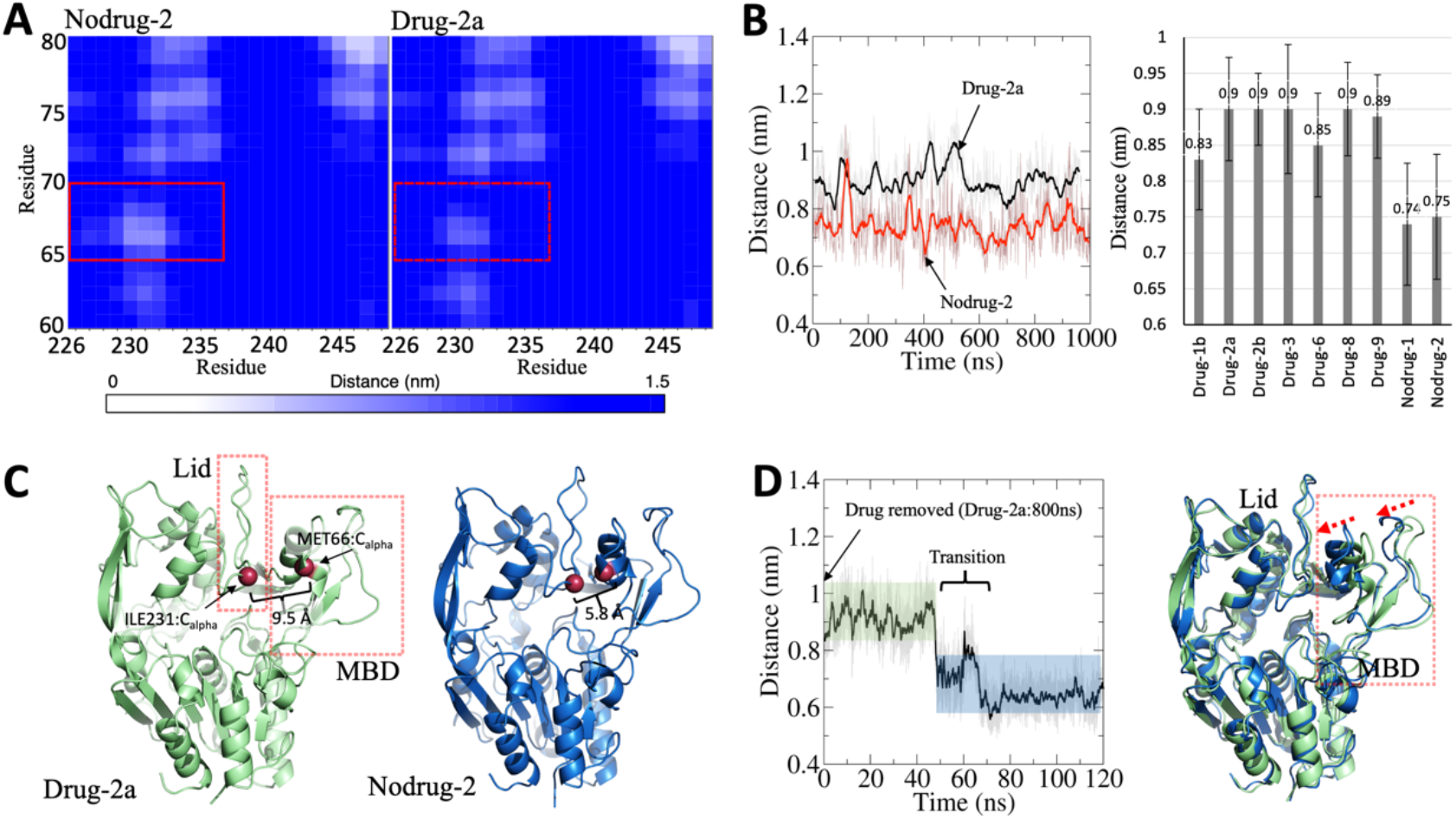
**(A)** The average distances between the relevant residues mapped as a distance matrix plot. Drug 2a is used as an example as the rest of the compounds induced a similar change. **(B)** The distance between the α-carbon atoms of M66 and I231 as a function of time in Drug-2a and Nodrug-2 simulations (Left). The average distances between the α-carbon atoms of M66 and I231 for all simulated systems (Right). **(C)** Snapshots from Drug-2a and Nodrug-2 simulation showing the maxima and minima of the detected conformational change. The proteins are rendered as cartoons and the marker residues’ α-carbons as red spheres. **(D)** The distance plot derived from the simulation where compound 2a was removed from the allosteric site after 800 ns (Left). Two simulation snapshots superimposed showing the orientation of the MBD domain before (0 ns; green) and after the removal of compound 2a (120 ns; blue) (Right).

We also determined the energetics of the MBD movement by utilizing the accelerated weight histogram method to calculate the free energy profiles as a function of the distance between the α-carbon atoms of M66 and I231. Free energy simulations were carried out with and without compound 2a. Fig 5 shows that the free energy minima at the distances of 0.7 and 0.9 nm agree with the distance analysis. Interestingly, the profiles show that when compound 2a is bound to the allosteric site, ~10 kJ/mol of energy is needed to move the MBD domain closer to the lid residing cavity. However, the energy required to move the MBD away from the lid cavity is smaller (~6 kJ/mol). This finding indicates that the drugs are hindering the MBD movement closer to the lid-residing cavity, which results in the broader cavity. Centered on this finding, we hypothesize that the wider cavity promotes the lid movement from the closed state to an open one and *vice versa* due to lessened steric hindrance. To shed more light on this, we transformed the lid loop from the open state to a closed state by slowly pulling the lid loop towards the active site tunnel opening. At the same time, we monitored the distance of the α-carbon atom of M66 from the initial position as a function of pulling time. As seen in Fig 6, the MBD domain moves away from the initial position (~0.3-0.5 nm) when the lid changes its conformation from an open to a closed state. However, we must remark that our pulling experiments do not necessarily resemble the correct conformational change pathway for the lid as the lid’s transitional pathway is unknown. Nevertheless, our pulling results further support the hypothesis that the MBD needs to shift its orientation to render the lid’s conformational shift possible.

**Figure 5.**
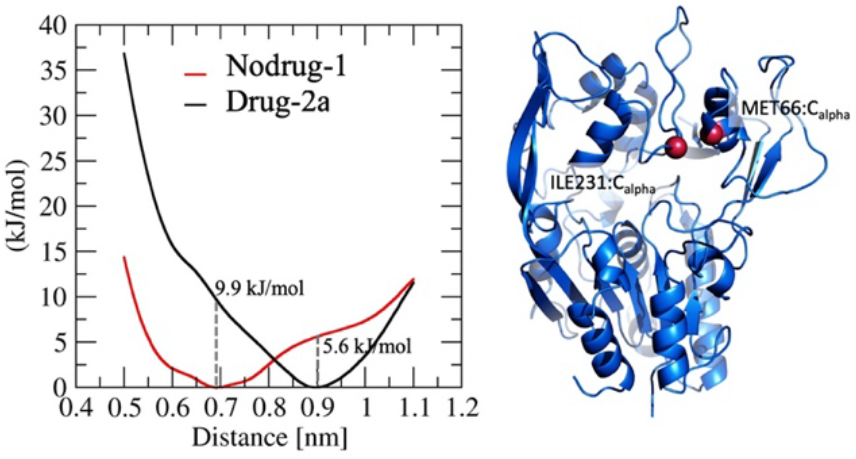
Spatial energetics of the MBD. The free energy profiles as a function of the distance between M66 and I231 α-carbon atoms with and without compound 2a (Left). A blue cartoon presentation of LCAT showing the ILE231 and MET66 Cα atoms (red spheres) that were used to determine the conformational free energy as a function of distance (Right).

**Figure 6.**
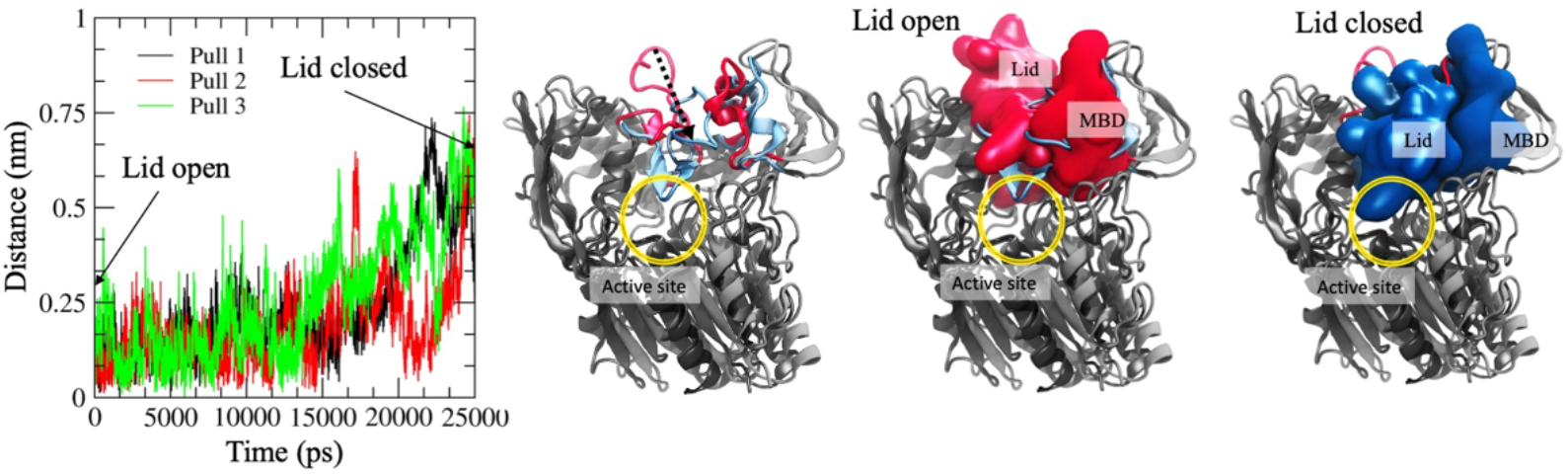
Steered molecular dynamics analysis showing the distance of the α-carbon atom of MET66 from the initial position as a function of time (Left). Superimposed snapshots illustrating the pulling pathway of the lid loop (Right). The light red and blue cartoon or van der Waals renderings mark the lid loop’s open and closed states, respectively. The rest of the LCAT enzyme is rendered as a grey cartoon representation. The black arrow indicates the movement of the lid loop during the pulling simulations. The active site tunnel opening is marked with a yellow sphere.

In the light of these findings, it is tempting to hypothesize that the MBD’s reangling is responsible for the increased activity of LCAT bound with different drugs in the MUP esterase assays reported by Manthei et al. (18). Intriguingly, while compound 6 induced the same orientational change, it did not promote LCAT activity in the experiments. This is likely due to the compound’s inability to bind to LCAT, as was shown by a microscale thermophoresis analysis conducted in the same study. In our docking calculations, compound 6 is directly docked to a favorable position and, thus, shows favorable binding free energies, which naturally does not consider the drug’s introduction into the allosteric site of LCAT. Namely, the entry of compound 6 may be accompanied by a large free energy barrier, which kinetically prevents the interaction with the allosteric site. Thus, our results imply that the pyrazine ring is required for the entry into the allosteric site of LCAT, but it is not critical when the activation of LCAT is mechanistically considered. Therefore, the variation of chemical groups in the pyrazine ring might provide the most robust means to tune the current set of positive allosteric modulators’ potency.

A speculative cause of this lack of binding could be the inability to participate in a favorable inter-protein interaction. A pseudo-2-fold interface was detected in the crystal structure, where the bound compounds’ tail rings formed a bridge to the opposing LCAT (18). This homodimer form could promote allosteric modulator binding by helping the compounds overcome the suggested free energy barrier. Although it was not possible to produce similar stable crystals with compounds 2 and 3, the pyrazine ring’s gained reach could have allowed a partial homodimerization where the compounds were locked in their places; the subsequently formed steric hindrance prevented the homodimer from stabilizing. Regardless, the pyrazine ring may also have a different role in HDL-binding, as a recent model of LCAT-HDL binding shows the tail of compound 1b poking out of the MBD (41), where it may affect the orientation of LCAT at the lipid-water interface.

More research needs to be done to elucidate the positive effect of the pyrazine ring on the entry into LCAT, which would further characterize an optimal allosteric modulator. Although we could not establish a clear connection between the compounds’ binding energies or induced effects and their experimental potencies, the conformational change discovered here provides a plausible explanation of the mechanism of the effect the compounds elicit in LCAT. This is significant as the observed conformational change can be used as a computational screening tool to discover and develop new positive allosteric modulators for LCAT, benefiting the pharmaceutical research on LCAT related diseases.

## CONCLUSIONS

In this research study, we aimed to characterize how a set of positive allosteric modulators interact with LCAT, firstly, to produce new insights on how these modulators mechanistically promote the activity and, secondly, shed light on the general activation mechanism of LCAT mediated by different apolipoproteins, chiefly apoA-I. While no apparent differences in binding or mode of interaction were registered between the allosteric modulators that could explain their different fold activities, we found out that all molecules can realign the MBD of LCAT. Further, our steered molecular dynamics simulations suggest that the MBD needs to retract away from the lid binding cavity during the lid’s conformational shift from the closed to the open state and *vice versa*. Thus, we hypothesize that the realignment and the altered free energy landscape of the MBD induced by positive allosteric modulators facilitate the activation of LCAT by making it easier for the lid loop to alternate between open and closed conformations. This is because the MBD’s realignment may either simply lessen the steric hindrances associated with the change or modulate the transitional folding pathway’s energetics. The findings presented in this study can be possibly validated by nuclear magnetic resonance studies or by producing crystal structures for LCAT with solely an allosteric compound bound to LCAT without IDFP. To sum up, our findings provide a plausible explanation of why the set of positive allosteric modulators studied here increases the fold rate of LCAT. This information can be exploited to design new LCAT activating drug molecules to treat LCAT deficiencies and CVDs or further understand the activation mechanism of LCAT mediated by apoA-I.

## ACKNOWLEDGEMENTS

Academy of Finland, grant, 298863 (AK), have provided financial support. All computational resources provided by the Finnish IT Centre for Science Ltd. (CSC).

## REFERENCES

1. Lewis GF, Rader DJ. New Insights Into the Regulation of HDL Metabolism and Reverse Cholesterol Transport. Circ Res. 2005 June 24;96(12):1221–32.

2. Haase CL, Tybjærg-Hansen A, Ali Qayyum A, Schou J, Nordestgaard BG, Frikke-Schmidt R. LCAT, HDL Cholesterol, and Ischemic Cardiovascular Disease: A Mendelian Randomization Study of HDL Cholesterol in 54,500 Individuals. J Clin Endocrinol Metab. 2012 Feb;97(2):E248–56.

3. Rader DJ, Hovingh GK. HDL and cardiovascular disease. Lancet. 2014 Aug 16;384(9943):618–25.

4. Peloso GM, Auer PL, Bis JC, Voorman A, Morrison AC, Stitziel NO, et al. Association of low-frequency and rare coding-sequence variants with blood lipids and coronary heart disease in 56,000 whites and blacks. Am J Hum Genet. 2014 Feb 6;94(2):223–32.

5. Carlson LA, Philipson B. Fish-eye disease. A new familial condition with massive corneal opacities and dyslipoproteinaemia. Lancet. 1979 Nov 3;2(8149):922–4.

6. Kuivenhoven JA, Pritchard H, Hill J, Frohlich J, Assmann G, Kastelein J. The molecular pathology of lecithin:cholesterol acyltransferase (LCAT) deficiency syndromes. J Lipid Res. 1997 Feb;38(2):191–205.

7. Kuivenhoven JA, Voorst, E. J. van Voorst tot, Wiebusch H, Marcovina SM, Funke H, Assmann G, et al. A unique genetic and biochemical presentation of fish-eye disease. J Clin Invest. 1995 Dec;96(6):2783–91.

8. Shamburek RD, Bakker-Arkema R, Shamburek AM, Freeman LA, Amar MJ, Auerbach B, et al. Safety and Tolerability of ACP-501, a Recombinant Human Lecithin: Cholesterol Acyltransferase, in a Phase 1 Single-Dose Escalation Study. Circ Res. 2016 Jan 8;118(1):73–82.

9. Gunawardane RN, Fordstrom P, Piper DE, Masterman S, Siu S, Liu D, et al. Agonistic Human Antibodies Binding to Lecithin-Cholesterol Acyltransferase Modulate High Density Lipoprotein Metabolism. J Biol Chem. 2016 Feb 5;291(6):2799–811.

10. Freeman LA, Demosky SJ, Konaklieva M, Kuskovsky R, Aponte A, Ossoli AF, et al. Lecithin:Cholesterol Acyltransferase Activation by Sulfhydryl-Reactive Small Molecules: Role of Cysteine-31. J Pharmacol Exp Ther. 2017 Aug;362(2):306–18.

11. Freeman LA, Karathanasis SK, Remaley AT. Novel lecithin: cholesterol acyltransferase-based therapeutic approaches. Curr Opin Lipidol. 2020 Apr;31(2):71–9.

12. A Phase 2 a Randomized, Double-blind, Placebo-controlled Study to Evaluate the Safety, Pharmacokinetics, and Pharmacodynamics of Single-Ascending Doses of MEDI6012 in Subjects With Stable Coronary Artery Disease. 2018 Mar 16.

13. Gerl MJ, Vaz WLC, Domingues N, Klose C, Surma MA, Sampaio JL, et al. Cholesterol is Inefficiently Converted to Cholesteryl Esters in the Blood of Cardiovascular Disease Patients. Sci Rep. 2018 Nov 3;8(1):1–11.

14. Kobayashi H, Ohkawa N, Takano M, Kubota H, Onoda T, Kaneko T, et al., inventors; Daiichi Sankyo Co Ltd, assignee. Piperidinylpyrazolopyridine derivative. United States patent US 9150575B2. 2015.

15. Kobayashi H, Ohkawa N, Takano M, Kubota H, Onoda T, Kaneko T, et al., inventors; Condensed pyrazole derivative. World Intellectual Property Organization WO 2015111545. 2015.

16. Onoda T, Kaneko T, Arai M, Kobayashi H, Terasaka N, inventors; Imidazopyridine derivative. World Intellectual Property Organization WO 2015087996. 2015.

17. Kobayashi H, Arai M, Kaneko T, Terasaka N, inventors; Daiichi Sankyo Co Ltd, assignee. 5-hydroxy-4-(trifluoromethyl)pyrazolopyridine derivative. United States patent US 9796709. 2016.

18. Manthei KA, Yang S, Baljinnyam B, Chang L, Glukhova A, Yuan W, et al. Molecular basis for activation of lecithin:cholesterol acyltransferase by a compound that increases HDL cholesterol. eLife. 2018 Nov 27;7:e41604.

19. Casteleijn MG, Parkkila P, Viitala T, Koivuniemi A. Interaction of lecithin:cholesterol acyltransferase with lipid surfaces and apolipoprotein A-I-derived peptides. J Lipid Res. 2018 Apr;59(4):670–83.

20. Peelman F, Vanloo B, Perez-Mendez O, Decout A, Verschelde J, Labeur C, et al. Characterization of functional residues in the interfacial recognition domain of lecithin cholesterol acyltransferase (LCAT). Protein Eng Des Sel. 1999 Jan;12(1):71–8.

21. Adimoolam S, Jonas A. Identification of a Domain of Lecithin–Cholesterol Acyltransferase That Is Involved in Interfacial Recognition. Biochem Bioph Res Co. 1997 Mar 27;232(3):783–7.

22. Jin L, Shieh J, Grabbe E, Adimoolam S, Durbin D, Jonas A. Surface Plasmon Resonance Biosensor Studies of Human Wild-Type and Mutant Lecithin Cholesterol Acyltransferase Interactions with Lipoproteins. Biochemistry. 1999 Nov;38(47):15659–65.

23. Manthei KA, Ahn J, Glukhova A, Yuan W, Larkin C, Manett TD, et al. A retractable lid in lecithin:cholesterol acyltransferase provides a structural mechanism for activation by apolipoprotein A-I. J Biol Chem. 2017 Dec 8;292(49):20313–27.

24. Webb B, Sali A. Protein structure modeling with MODELLER. Methods Mol Biol. 2014;1137:1–15.

25. Hanwell M, Curtis D, Lonie DC, Vandermeersch T, Zurek E, Hutchison G. Avogadro: an advanced semantic chemical editor, visualization, and analysis platform. J Cheminformatics. 2012;4(17).

26. Trott O, Olson AJ. AutoDock Vina: improving the speed and accuracy of docking with a new scoring function, efficient optimization and multithreading. J Comput Chem. 2010 Jan 30;31(2):455–61.

27. Lindorff-Larsen K, Piana S, Palmo K, Maragakis P, Klepeis JL, Dror RO, et al. Improved side-chain torsion potentials for the Amber ff99SB protein force field. Proteins, structure, function, and bioinformatics. Proteins. 2010 Jun;78(8):1950–8.

28. Jorgensen WL, Chandrasekhar J, Madura JD, Impey RW, Klein ML. Comparison of simple potential functions for simulating liquid water. J Chem Phys. 1983 Jul 15;79(2):926–35.

29. Frisch MJ, Trucks GW, Schlegel HB, Scuseria GE, Robb MA, Cheeseman JR, et al. Gaussian 16 Revision A.03. 2016.

30. Singh UC, Kollman PA. An approach to computing electrostatic charges for molecules. J Comput Chem. 1984;5(2):129–45.

31. Case DA, Ben-Shalom IY, Brozell SR, Cerutti DS, Cheatham III TE, Cruzeiro VWD, et al. AMBER 2018. 2018.

32. Wang J, Wolf RM, Caldwell JW, Kollman PA, Case DA. Development and testing of a general amber force field. J Comput Chem. 2004 Jul 15;25(9):1157–74.

33. Bussi G, Donadio D, Parrinello M. Canonical sampling through velocity-rescaling. J Chem Phys. 2008 Mar 28.

34. Parrinello M, Rahman A. Polymorphic transitions in single crystals: A new molecular dynamics method. J Appl Phys. 1981 Dec 1;52(12):7182–90.

35. Darden T, York D, Pedersen L. Particle mesh Ewald: An N⋅log(N) method for Ewald sums in large systems. J Chem Phys. 1993 Jun 15;98(12):10089–92.

36. Berendsen HJC, van der Spoel D, van Drunen R. GROMACS: A message-passing parallel molecular dynamics implementation. Comput Phys Commun. 1995 Sep 2;91(1):43–56.

37. Abraham MJ, Murtola T, Schulz R, Páll S, Smith JC, Hess B, et al. GROMACS: High performance molecular simulations through multi-level parallelism from laptops to supercomputers. SoftwareX. 2015 Sep 1;1-2:19-25.

38. Lindahl V, Lidmar J, Hess B. Accelerated weight histogram method for exploring free energy landscapes. J Chem Phys. 2014 Jul 24;141(4):044110.

39. Lidmar J. Improving the efficiency of extended ensemble simulations: The accelerated weight histogram method. Phys Rev E. 2012 May 25;85(5):056708.

40. Humphrey W, Dalke A, Schulten K. VMD: visual molecular dynamics. J Mol Graph. 1996 Feb;14(1):33–38.

41. Manthei KA, Patra D, Wilson CJ, Fawaz MV, Piersimoni L, Shenkar JC, et al. Structural analysis of lecithin:cholesterol acyltransferase bound to high density lipoprotein particles. Commun Biol. 2020 Jan 15;3(1):1–11.

